# Reversing Eroom’s Law: Finitude and Dilution in Small-Molecule Discovery

**DOI:** 10.1101/2025.10.29.685434

**Authors:** Ross Youngs

**Affiliations:** Biosortia, Inc., 2545 Farmers Dr., Suite 370, Columbus, OH 43235

## Abstract

Drug discovery productivity—measured as clinically viable new molecules per research has declined for decades, a trend known as Eroom’s Law. This paradox persists despite advances in screening, computing, and artificial intelligence. Here we show that it stems from two forces: the finite diversity of nature’s biosynthetic rules, leading to saturation of novel natural scaffolds, and the dilution of viable candidates in expanding synthetic chemical spaces, where success rates fall without improved specificity. We propose an optimized hybrid strategy that allocates effort to mining natural products and using their validated patterns to guide synthetic exploration, maximizing novelty per cost. This approach yields testable rules, such as optimal allocation thresholds and benchmarks for artificial intelligence tools. By embedding natural principles in modern discovery, this framework offers a path to reverse Eroom’s Law and boost pharmaceutical efficiency.

## Introduction: The paradox and the thesis

Across the pharmaceutical industry, approvals per inflation-adjusted research and development dollar have fallen even as screening, synthesis and computing have expanded,^1–4^ a pattern termed Eroom’s law. Here we advance a mechanistic explanation with three coupled results and a practical remedy. The framework rests on three ideas: nature’s finite biosynthetic playbook explains diminishing returns in natural-product discovery; the dilution of synthetic design space explains analogous declines in human-designed exploration; and a finite-prior hybrid that leverages natural priors to steer synthetic search provides a route to recover productivity. Empirical anchors include enumerations of chemical space,^5–7^ clinical success rates and analyses of research-and-development efficiency.^1–4^ Throughout, we use “discovery/returns” to mean clinically viable molecular novelty per unit of cost (proxied by approvals or late-stage successes per inflation-adjusted research and development dollar) rather than the number of molecules screened. Formal notation and parameter definitions are provided in Box 1 and the Methods to keep the Introduction accessible to a broad readership.

### Nature’s finite playbook and diminishing molecular returns

The saturation of *N* discoverable classes over cumulative effort Σ can be modeled as a finite-class arrival process:

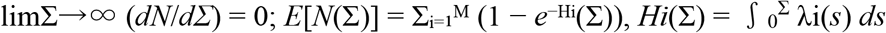

Across geological time, biochemistry has used a finite element palette with stability/complexity windows and only a finite number of biosynthetic grammar expansions. Upon broad exploration, the accessible set *A*\* is effectively finite and class-level novelty saturates as effort increases. First-arrival hazards over *M* classes yield 1 − *e*−*Hi* with *Hi* →∞ and bounded hazards, and dominated convergence gives *E* [*N*] → *M* and *dE* [*N*]/*dΣ* → 0. Standard arguments imply almost-sure saturation. Punctuations — rare grammar additions—add finite increments to *M*, producing transient spikes before decline resumes.^8,9^ Modeled as a non-homogeneous Poisson process with decreasing intensity μ(Σ), punctuations add Poisson(ν) class increments at times T_k_, causing dN/dΣ spikes (≈ ν λ_avg_) with lengthening intervals τ_k_, testable via taxonomic databases. While total biospheric variants are vast, most new finds in unexplored taxa and cryptic clusters are variants on the existing scaffolds generated by a finite grammar; hence, class discovery faces diminishing returns even as variants abound. Large resources (e.g., COCONUT) show redundancies consistent with this view.^10–12^ Moreover, many plant-derived molecules are endophyte encoded or governed by epigenetic on/off states— evidence of a shared grammar *being leveraged across different organisms* rather than open-ended class creation.^13^

**Figure 1: Saturation *N*(*Σ*) and rate *dN*/*dΣ* with dashed lines at punctuation events**

**Figure 1.**
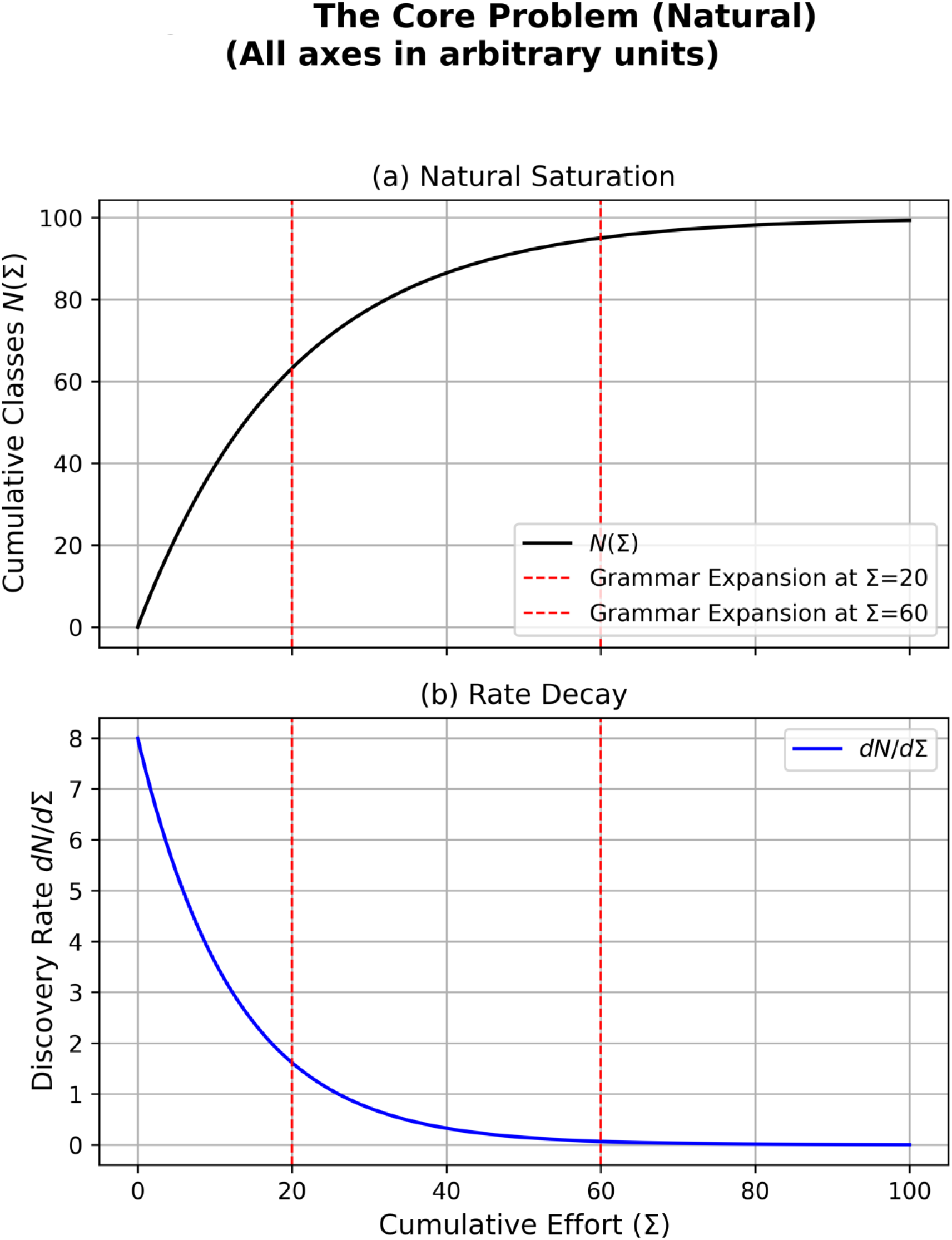
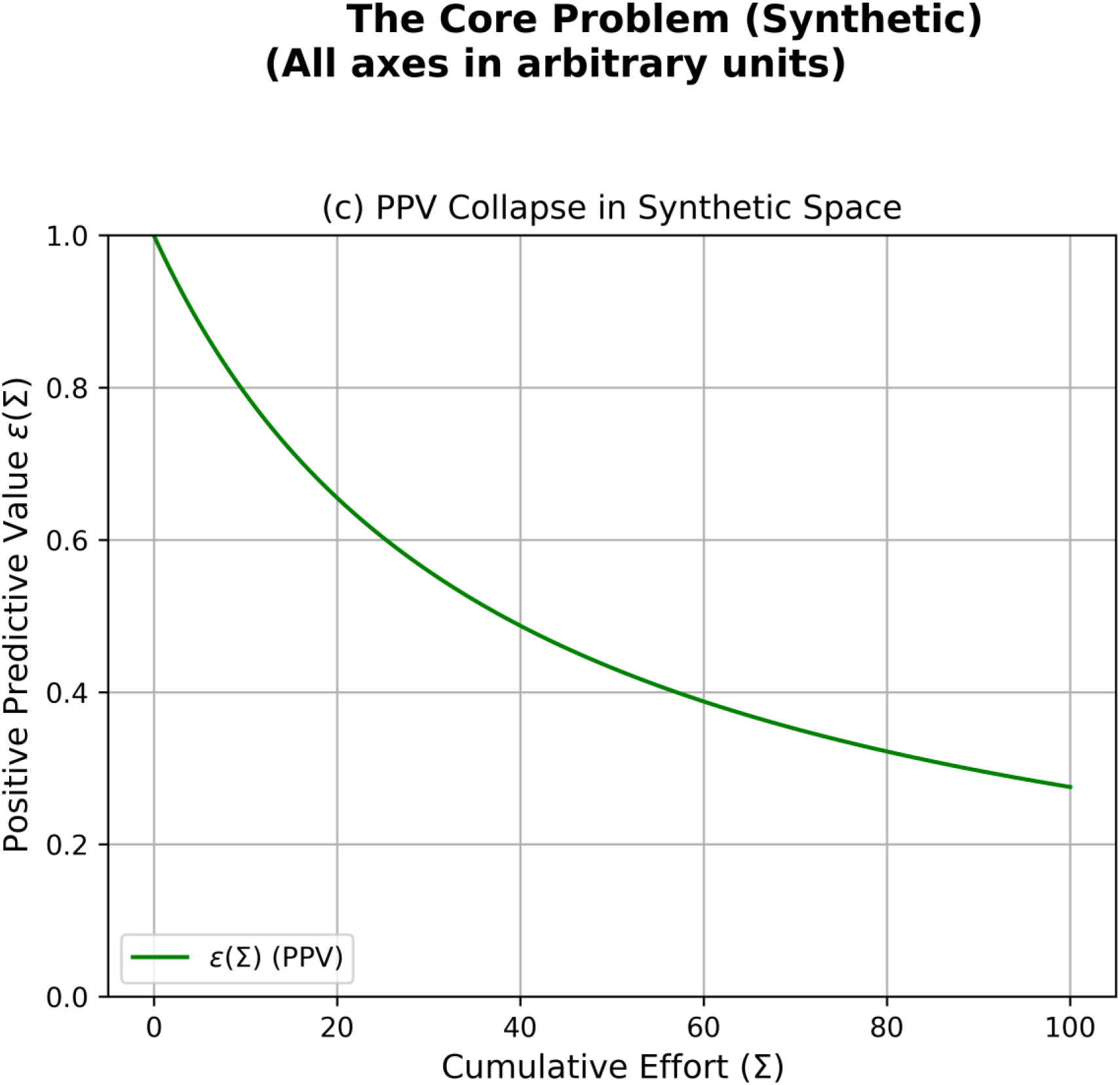
The core problem. (a) Natural saturation: *N(Σ)* saturates with rare grammar expansions (vertical dashed lines). (b) Rate decay: *dN/dΣ* exhibits transient spikes at expansions and decays toward zero. (c) PPV collapse: *ε(Σ)* falls as *π(Σ)* declines; modest *β(Σ)* gains are insufficient.

### Infinite-horizon dilution in synthetic space

Let *U*(*Σ*) be the design universe with |*U*| → ∞,^5–7^ including synthesizable vendor libraries (e.g., Enamine REAL Space).^14^ Let *V* be truly viable classes with prevalence *π*(*Σ*) = |*V*∩*U*|/|*U*|. Viability requires satisfying multiple orthogonal constraints (efficacy, selectivity, safety, and ADME), so *π*(*Σ*) → 0 as the universe expands. Pipelines with sensitivity *α*(*Σ*) and false-positive rate *β*(*Σ*) yield positive predictive value (PPV) *ε*(*Σ*) = *α*/[*α*+*β*(1−*π*)/*π*]. As *π* → 0, *ε* collapses unless specificity improves commensurately; with finite enrichment and throughput, the viable-discovery rate *Λε* falls toward zero.^15–16^

**Figure 1c: PPV collapse *ε*(*Σ*) vs. *π*(*Σ*)**

**Figure 2: Sublinear growth of *N*(*Σ*) and decline of *dN*/*dΣ* under polynomial case**

**Figure 2.**
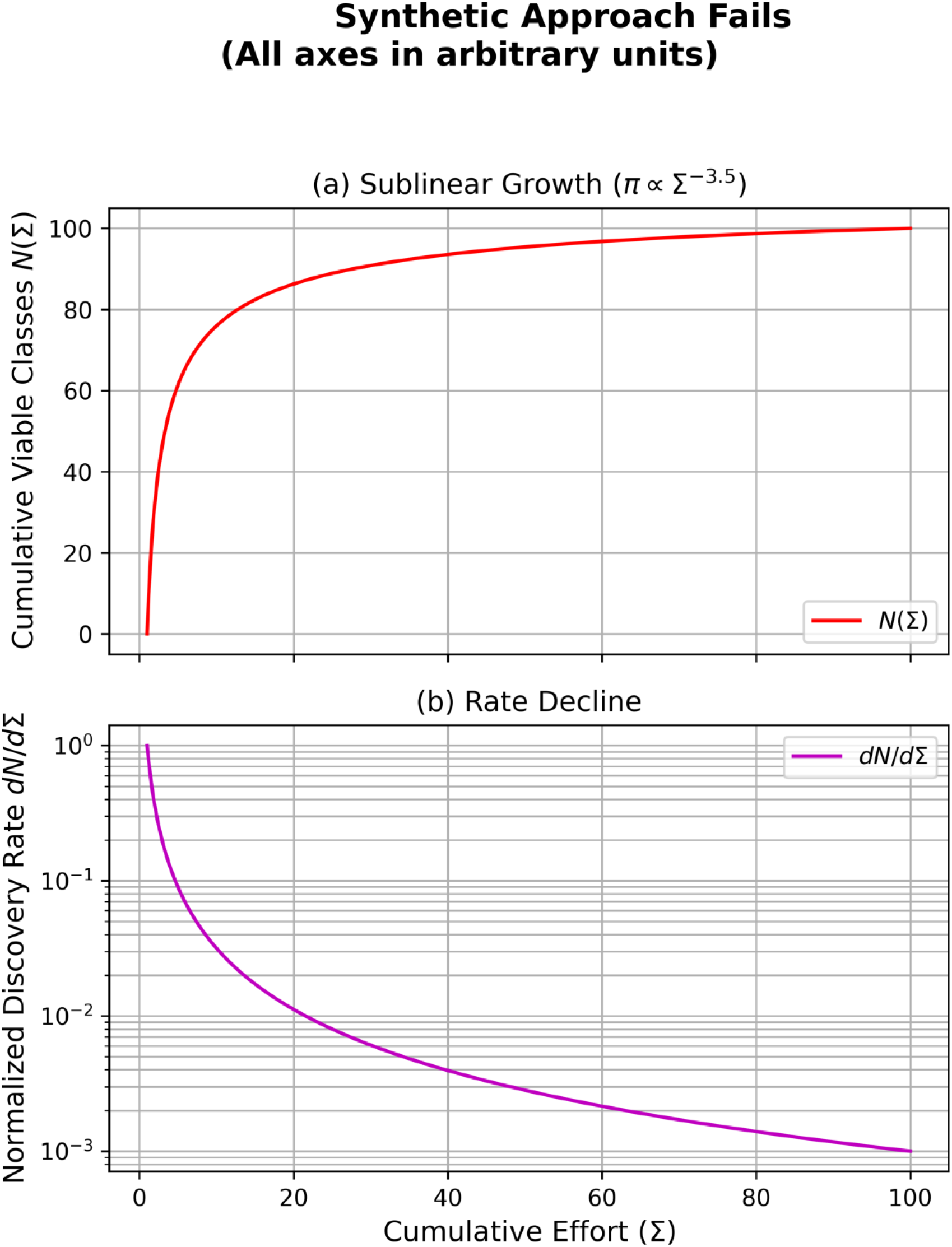
Synthetic approach fails without super-linear enrichment. (a) Sublinear growth: Cumulative viable classes *N(Σ)* grow sublinearly when π(Σ) ∝ Σ^−(k+1+η)^. (b) Rate decline: *dN/dΣ* declines with *Σ* under the same parameters.

Polynomial sufficiency: If *Λ*(*Σ*) ∝ *Σ*^*k*^ but π(*Σ*) ∝ *Σ*^−(*k*+1+*η*)^ (for some *η*> 0), then *Λε* → 0.

Exponential counter—case (AI): If *π* ≈ *e*^−*rΣ*^ and *Λ* ≈ *e*^*cΣ*^, then *Λε* ≈ (*α*/*β*) *e*^(*c*−*r*)*Σ*^, and dilution is mitigated only if *c* > *r* while *β* falls—which provides a testable AI benchmark for current models.^16^ Public repositories continue to document extreme scale growth (e.g., PubChem 2025 update),^17^ and recent AI-agent pipelines illustrate how throughput may scale operationally.^18^

### Hybrid strategy with data dependency

The hybrid model’s viable-discovery rate and its return-per-cost, ℛ are defined by the following relationships:

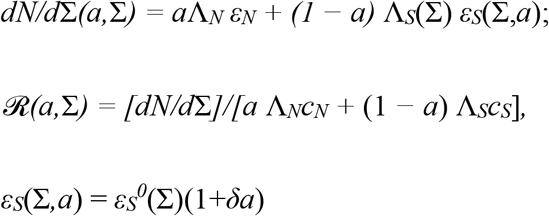

Allocate a share *a* ∈ [0,1] to the finite/natural channel (*N*) and 1−*a* to the expanding/synthetic channel (*S*). The hybrid’s viable-discovery rate and return-per-cost *are governed by* data dependency (δ): The transfer of natural priors into *S* is captured as a multiplicative PPV lift εS (Σ, *a*) = *ε*_*S*_^0^ (Σ)[1+δ*a*]. When δ > 0, investing in *N* improves the PPV of *S*; when δ =0, channels are decoupled. Equal-marginal policy gate (breakeven): shift marginal effort from *N* to *S* only if *ε*_*S*_^0^ (Σ)[1+δ*a*] ≥ (*cS* / *cN*) εN. With data dependency, this yields a closed-form lower bound for interior optima: *a*\* (Σ) ≥ [(*cS* / *cN*) εN / *ε*_*S*_^0^ (Σ) −1] / δ (clipped to [0,1]). As prevalence in *S* collapses (*ε*_*S*_^0^↓), *a*\* rises; larger δ lowers the breakeven PPV that *S* must achieve.

**Figure 3: Shifting optimum *a*\*(*Σ*) and bar chart showing hybrid dominating pure strategies**

**Figure 3.**
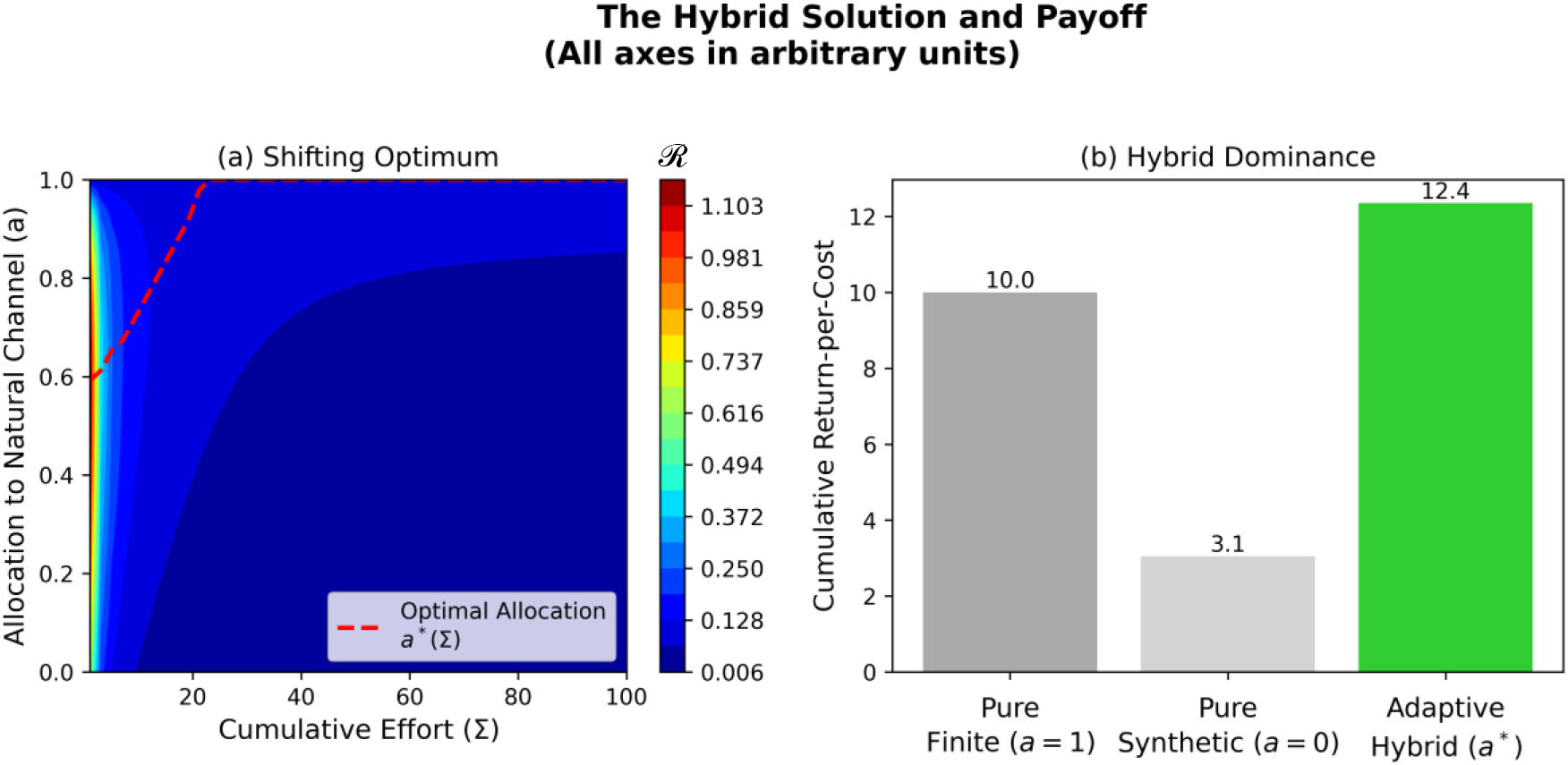
The hybrid solution and payoff. (a) Shifting optimum: Return-per-cost *ℛ(a,Σ)* exhibits interior optima that shift toward larger a as Σ grows and *ε*_S_^0^ falls; larger δ lowers breakeven PPV. (b) Dominance: Cumulative return-per-cost—hybrid (adaptive *a\**) vs. pure finite (a = 1) vs. pure infinite (a = 0). Bars were plotted on a scientific y-axis; labels were omitted to avoid misleading zeros.

### Falsifiable predictions

P1 (allocation gate): At a decision horizon *Σ*_*ref*_, if *ε*_*S*_^0^_*S* (*Σ_ref*)_< (*c*_*S/c_N*_) *ε*_*N*_ / (1+*δa*), programs do not shift to *S*; they invest in *N* or reduce *β* until above gate.

P2 (AI benchmark): Exponential counter—case is passed only if measured *c* ≥ *r* and *β(Σ)* declines—falsifiable in prospective pipelines.^16,18^

P3 (punctuations): Natural-channel dN/dΣ exhibits transient spikes post-punctuation (modeled on PAGE 3), with lengthening intervals τ_k_, falsifiable via taxonomic cluster analysis. ^8,9^

P4 (reverse-Eroom regime): If (1+*δa*) > (*c*_*S*/*cN*)_ (*ε*_*N*/*εS*_^0^) (with defaults δ ≈ 1, a ≈ 0.5, c_S_ / c_N_ ≈ 1, ε_N_ / *ε*_*S*_^0^ ≈ 1.5), a hybrid increases novelty per cost versus pure *S* and can raise approvals per dollar; this is testable in portfolio trials.

## Discussion and validation paths

This framework explains finite closure in nature, base-rate dilution in synthetic space and why a non-zero allocation to natural priors is necessary for sustainable productivity. It reframes Eroom’s law as a predictable consequence of saturation and dilution, rather than a failure of technology, and indicates that the long-standing emphasis on synthetic discovery and the resultant dilution becomes unsustainable at scale. Reversing this trend requires a portfolio shift toward a hybrid model, in which finite-prior discovery improves the efficiency and specificity of synthetic and artificial intelligence (AI) pipelines. Validation can proceed along three fronts. First, redundancy analyses in large natural-product corpora to quantify late-stage saturation at the class level even as variants continue to grow.^10–12^ Second, prospective measurements in modern screening to track the base rate (prevalence), sensitivity and false-positive rate, and to test both the allocation gate and an AI benchmark in which throughput growth outpaces prevalence decay while the false-positive rate falls.^5–7,13–18^ Third, portfolio-level trials of the equal-marginal decision rule—allocating effort until marginal return per cost equalizes across channels—with preregistered endpoints of novelty per cost and late-stage success. This perspective also reorients goals for AI and synthetic biology: not only to enumerate molecules in unexplored niches, but to learn and strategically expand nature’s finite biosynthetic grammar, including endophyte and lichen consortia, through targeted epigenetic and pathway engineering under finite-prior constraints.^14^ We account for problem depth by using a difficulty-adjusted positive predictive value (see Box 1), which makes explicit that deeper problems require stronger finite-prior lift, higher specificity, greater throughput or lower unit cost to sustain returns.

### Box 1 | Formal framework

This box defines symbols and conventions used in the framework and appendix. Nonstandard symbols are defined here. Indices are shown as subscripts and exponents as superscripts.

### Sets and spaces

- *U* — design universe (set of all classes/constructs considered)
- *V* — viable subset of *U* (those meeting the criteria below)

### Functions and quantities

- *Σ* — cumulative discovery/effort variable used as the independent axis
- *π(Σ)* — prevalence of viable classes (fraction of *U* that is viable). As |*U*| → ∞, *π(Σ)*→ 0
- *λ*_*i*_*(Σ)* — instantaneous hazard/intensity for class *i* at cumulative effort *Σ*
- *H*_*i*_*(Σ)* — cumulative hazard for class *i*. For example, the survival term uses e^−Hi(Σ)^
- *Λ(Σ)* — throughput/attempt rate as a function of *Σ*
- *ε(Σ)* — PPV under the screening pipeline

### Indices, strategies, and constants

- *i, j, k* — class indices (subscripts)
- *N, S* — strategy tags (narrow *N*; sweep *S*), used as subscripts (e.g., *Λ*_*N*_, *ε*_*S*_)
- *ref* — reference point tag, used as a subscript (e.g., *Σ*_*ref*_)
- *δ, α, β, r, c* — constants/parameters as defined in context (screening sensitivity *α*, false-positive rate *β*; exponents/rates *r, c*; response slope *δ*)
- *ε*_*S*_^0^ — baseline PPV for the sweep strategy (*S*), used in response models

### Operators and notation

- **Σ**_i=1_^M^ — summation over classes i = 1, …, M
- ∫_0_^Σ^ (…) ds — integral from 0 to *Σ* with respect to *s*
- *→*, ≤, *≥, ≈* — limit, inequality, and approximation symbols as standard

### Conventions

- Indices appear as subscripts (e.g., *λ*_*i*_, *H*_*i*_, *ε*_*S*_), and exponents appear as superscripts (e.g., Σ^k^).
- Exponentials are written as e^n^ with the entire exponent in a single superscript run (e.g., e^−Hi(Σ)^).
- Greek and special symbols (*Σ, Λ, π*, →, ≤, ≥) are preserved; prose and punctuation follow journal style.
- Default parameters for predictions (e.g., P4): δ ≈ 1 (moderate prior lift), a ≈ 0.5 (balanced allocation), c_S_ / c_N_ ≈ 1 (equal costs), ε_N_ / *ε*_*S*_^0^ ≈ 1.5 (natural PPV advantage),, and γ ∈ [0,1], default γ = 1, for sublinear to linear complexity scaling in ε_eff_ = ε / ρ^γ^.

## Methods (summary)

### Natural

Finite-class first-arrival process; dominated/monotone convergence; rare grammar shocks add finite increments^8–9^

### Synthetic

PPV/base-rate calculus; *π(Σ)* decays with *Λ(Σ)* growth; sufficient polynomial and exponential conditions for *Λε* → 0^5–7,14–16^

### Hybrid

Maximize *ℛ(a,Σ)*; equal-marginal rule; data dependency *δ* as PPV lift; sensitivity: ∂a*/∂*δ*> 0, ∂a*/∂(c_N_/c_S_)< 0

### Hybrid (difficulty extension)

We model task difficulty with *ρ* ≥ 1 and set ε_eff_ = ε/*ρ*^γ^ (γ∈ [0,1], default γ = 1); the equal-marginal gate and *a*^*^*(Σ)* follow by substitution.

## Data and code availability

No new data were generated in this study. The code for reproducing figures and the spreadsheet model are available upon reasonable request.

## Acknowledgements

The author thanks colleagues for feedback that strengthened the clarity, empirical grounding, and policy relevance of this work, with special thanks to Dr. Jack Scannell; Dr. David Newman, retired from NIH; and Professor Jonathan Eisen for their formative discussions. Additional thanks go to Professor Matt Bertin, Dr. Guy Carter, Professor Bill Gerwick, Professor Pieter Dorrestein, Dr. Colin Kruse, Dr. Martin Latterich, Dr. Ron Moss, Eugene Francis, and Rosemarie Trueman for a decade of engagement and feedback that helped bring this work to fruition.

## Author contributions

Ross Youngs conceived the theorems and wrote the manuscript.

## Main References

1. BIO / Informa / QLS. Clinical development success rates 2011–2020 (2021). https://go.bio.org/rs/490-EHZ-999/images/ClinicalDevelopmentSuccessRates2011_2020.pdf

2. Scannell, J. W. et al. Diagnosing the decline in pharmaceutical R&D efficiency. Nat. Rev. Drug Discov. 11, 191–200 (2012). 10.1038/nrd3681

3. Wouters, O. J., McKee, M. & Luyten, J. Estimated research and development investment needed to bring a new medicine to the market, 2009–2018. JAMA 323, 844–853 (2020). 10.1001/jama.2020.1166

4. Sertkaya, A. et al. Costs of drug development and research and development returns. JAMA Netw. Open 7, e2415445 (2024). 10.1001/jamanetworkopen.2024.15445

5. Ruddigkeit, L. et al. Enumeration of 166 billion organic small molecules in GDB-17. J. Chem. Inf. Model. 52, 2864–2875 (2012). 10.1021/ci300415d

6. Reymond, J.-L. The chemical space project. Acc. Chem. Res. 48, 722–730 (2015). 10.1021/ar500432k

7. Polishchuk, P. G. et al. Estimation of the size of drug-like chemical space based on GDB-17 data. J. Comput.-Aided Mol. Des. 27, 675–679 (2013). 10.1007/s10822-013-9672-4

8. O’Hagan, D. et al. Biosynthesis of an organofluorine molecule. Nature 416, 279 (2002). 10.1038/416279a

9. Pearson, A. et al. Sterol synthesis in the last eukaryotic common ancestor. Proc. Natl Acad. Sci. USA 100, 3707–3712 (2003). 10.1073/pnas.2536559100

10. Pye, C. R. et al. Retrospective analysis of natural products provides insights for future discovery trends. Proc. Natl Acad. Sci. USA 114, 5601–5606 (2017). 10.1073/pnas.1614680114

11. Navarro-Muñoz, J. C. et al. A computational framework to explore large-scale biosynthetic diversity. Nat. Chem. Biol. 16, 60–68 (2020). 10.1038/s41589-019-0400-9

12. Sorokina, M. & Steinbeck, C. COCONUT online. J. Cheminform. 13, 2 (2021). 10.1186/s13321-020-00478-9

13. Liu, R. et al. Unlocking the metabolic potential of endophytic fungi through epigenetics: a paradigm shift for natural product discovery and plant–microbe interactions. Nat. Prod. Rep. 42, 1690 (2025). 10.1039/d5np00028a

14. Enamine. REAL Space Navigator (accessed 14 Sept 2025). https://enamine.net/compound-collections/real-compounds/real-space-navigator

15. Ioannidis, J. P. A. Why most published research findings are false. PLoS Med. 2, e124 (2005). 10.1371/journal.pmed.0020124

16. Abramson, J. et al. Accurate structure prediction of biomolecular interactions with AlphaFold 3. Nature 630, 493–500 (2024). 10.1038/s41586-024-07487-w

17. Kim, S. et al. PubChem 2025 update. Nucleic Acids Res. 53, D1516–D1525 (2025). 10.1093/nar/gkae1059

18. Gao, S. et al. Empowering biomedical discovery with AI agents. Cell 187, 4881–4899 (2024). 10.1016/j.cell.2024.08.045

