## Supplementary Appendix for "Reversing Eroom’s Law: Finitude and Dilution in Small-Molecule Discovery"

**Supplementary Appendix 1**

**Policy / rates / definitions:**

**N(Σ), dN/dΣ:** Cumulative viable classes and instantaneous viable-discovery rate

**a, δ:** Allocation to finite channel (0–1) and data-dependency parameter for prior transfer into *S*; $\varepsilon_S(\Sigma, a) = \varepsilon_S^0(\Sigma)[1+\delta a]$

**(a, Σ):** novelty-per-cost = $[a\ \Lambda_N\ \varepsilon_N + (1-a)\ \Lambda_S\ \varepsilon_S] / [a\ \Lambda_N\ c_N + (1-a)\ \Lambda_S\ c_S]$

***ρ*** (problem depth): Task difficulty / constraint dimensionality; $\rho \geq 1$ (larger $\rho$ = deeper problem)

**Policy gate:** Shift to *S* only if $\varepsilon_S^0(\Sigma)(1+\delta a) \geq (c_S / c_N)\ \varepsilon_N$

**Difficulty-aware policy gate:** $\varepsilon_S^0\ (\Sigma)\ [1+\delta a]/\rho^\gamma \geq (c_S / c_N)\ \varepsilon_N$ (default $\gamma = 1$)

**Sufficient conditions:** Polynomial ($\Lambda \propto \Sigma^k$, $\pi \propto \Sigma^{-(k+1+\eta)} \Rightarrow \Lambda\varepsilon \to 0$); exponential counter—case ($\Lambda \approx e^c\Sigma$, $\pi \approx e^{-r}\Sigma \Rightarrow \Lambda\varepsilon \approx (\alpha/\beta)\ e^{(c-r)}\Sigma$)

**ε(Σ) (PPV):** $\alpha / (\alpha + \beta \cdot (1-\pi)/\pi)$, with sensitivity α and false-positive rate β

**Clarifying definitions (local):**

**Σ:** Cumulative discovery effort (independent axis)

**Λ(Σ):** Attempt/throughput rate as a function of *Σ*; $\Lambda_N$, $\Lambda_S$ denote narrow/sweep rates

**$c_N$, $c_S$**: Unit costs for narrow and sweep strategies, respectively

**$\varepsilon_S^0$**: Baseline PPV for the sweep strategy; $\varepsilon_S(\Sigma, a) = \varepsilon_S^0(\Sigma)[1+\delta a]$

**U**: The design universe, representing the set of all molecular classes under consideration

**V**: The subset of *U* containing all truly **viable** classes that meet the necessary criteria for success

**$\mathscr{R}$(a,Σ)**: The **return per cost** (or novelty per cost) of a given allocation strategy.

**N, S**: Subscripts used as strategy tags to distinguish between the **natural** (finite) and **synthetic** (expanding) discovery channels

**η (eta)**: A parameter in the polynomial case that helps define how quickly the prevalence of viable molecules decays

**ref**: A subscript used to denote a specific **reference point** or decision horizon, such as $\boldsymbol{\Sigma_{ref}}$

**|U|**: The **cardinality** or size of the set $U$

∩: The symbol for **set intersection**, so $|V \cap U|$ represents the number of elements common to both set $V$ and set $U$

**∝**: The symbol for **proportionality**, indicating that two quantities scale by a constant factor

$\partial a / \partial \delta^*$: The **partial derivative**, which measures the sensitivity or rate of change of the optimal allocation ($a^*$) with respect to the data-dependency parameter ($\delta$)

**$\lambda_i(\Sigma)$**: The **instantaneous hazard**; the rate of discovery for class $i$ at a specific level of effort $\Sigma$

**$H_i(\Sigma)$**: The **cumulative hazard**; the total accumulated "risk" of discovery for class $i$ up to effort $\Sigma$, calculated as the integral of the instantaneous hazard

**A. Natural closure (finite first-arrival process)**

Assume $M$ classes are indexed by $i$ with piecewise-continuous bounded hazards $\lambda_i(\Sigma) \geq 0$ under broad exploration.

Integrated hazard $H_i(\Sigma) = \int_0^\Sigma \lambda_i(s)\, ds$: First-arrival probability by $\Sigma$ is $1 - e^{-H_i(\Sigma)}$. Hence, $E[N(\Sigma)] = \Sigma_{i=1}^M (1 - e^{-H_i(\Sigma)}$. If $H_i(\Sigma) \to \infty$ for all $i$ in support and hazards are bounded, dominated convergence gives $E[N(\Sigma)] \to M$ and $dE[N]/d\Sigma = \sum i\, \lambda_i\, e^{-H_i} \to 0$; standard first-arrival arguments imply almost-sure convergence for $N(\Sigma)$. M is finite, with each $\lambda_i(\Sigma) > 0$ on some unbounded interval and bounded above, ensuring $H_i(\Sigma) \to \infty$ a.s., so $dE[N]/d\Sigma \to 0$ via dominated convergence, consistent with a finite playbook.

Rare grammar expansions add finite increments to $M$, yielding transient spikes.

**B. Infinite-horizon dilution (base-rate logic)**

Let $\pi(\Sigma) = |V \cap U|/|U|$ be viability prevalence with $\pi(\Sigma) \to 0$ as $|U| \to \infty$.

With sensitivity $\alpha$ and false-positive rate $\beta$, PPV $\varepsilon(\Sigma) = \alpha/[\alpha+\beta(1-\pi)/\pi]$. For small $\pi$, $\varepsilon \approx (\alpha/\beta)\cdot\pi$. Throughput $\mathit{\Lambda}(\mathit{\Sigma})$ yields viable-discovery rate $\mathit{\Lambda\varepsilon}$.

Sufficient conditions: (i) $\Lambda(\Sigma)\propto\Sigma^{k}$, $\pi(\Sigma)\propto\Sigma^{-(k+1+\eta)} \Rightarrow \Lambda\varepsilon\to 0$; (ii) $\Lambda \approx e^{c}\Sigma$, $\pi \approx e^{-r}\Sigma \Rightarrow \Lambda\varepsilon \approx (\alpha/\beta)e^{(c-r)}\Sigma$ (requires $c > r$ with declining $\beta$ to avoid dilution)

**C. Data-dependent hybrid and equal-marginal gate**

Allocate $a$ to $N$ and $1-a$ to $S$. Let $\varepsilon_s(\Sigma,a) = \varepsilon_s^0(\Sigma)[1+\delta a]$ ($\delta\geq 0$). Rate: $dN/d\Sigma = a\Lambda_n\varepsilon_n + (1-a)\Lambda_s\varepsilon_s(\Sigma,a)$.

Return-per-cost: $\mathscr{R}$ = Rate / $[a\Lambda_n c_n + (1-a)\Lambda_s c_s]$

Equal-marginal rule $\Sigma_{re}\boldsymbol{f}$: Shift to $S$ if $\varepsilon_s^0(\Sigma_{re}\boldsymbol{f})(1+\delta a) \geq (c_s/c_n)\,\varepsilon_n$

Lower bound for interior optimum: $a^*(\Sigma_{re}\boldsymbol{f}) \geq \{[(c_s/c_n)\,\varepsilon_n / \varepsilon_s^0(\Sigma_{re}\boldsymbol{f})] - 1\} / \delta$ (clipped to [0,1]) Sensitivities: $\partial a^*/\partial\delta>0$; $\partial a^*/\partial\delta > 0$; $\partial a^*/\partial(c_n/c_s) < 0$; as $\varepsilon_s^0$ decreases with $\mathit{\Sigma}$, $a^*$ increases.

Notes: Symbols as defined in Box 1

**Supplementary Appendix 2**

**Full Proofs for the Falsifiable Predictions (P1–P4)**

The paper "Reversing Eroom's Law" outlines four quantitative, falsifiable predictions (P1–P4) derived from its unified framework of natural finitude, synthetic dilution, and finite-prior hybrid strategies. These predictions are grounded in the mathematical models presented in the appendix (Sections A–C). Below, I derive full proofs for each, expanding the compact versions in the appendix with step-by-step reasoning, explicit assumptions, and derivations. I use the paper's notation consistently:

- $\Sigma$: Cumulative discovery effort (proxy for cost/time/compute).
- $N(\Sigma)$: Cumulative viable molecular classes discovered.
- $dN/d\Sigma$: Instantaneous viable-discovery rate.
- $\Lambda(\Sigma)$: Throughput/attempt rate (e.g., molecules screened per unit $\Sigma$); subscripts N (natural/finite channel) and S (synthetic/expanding channel).
- $c_N$, $c_S$: Unit costs for natural and synthetic strategies.
- $\varepsilon$: Positive predictive value (PPV; fraction of screened candidates that are viable).
- $\pi(\Sigma)$: Prevalence of viable candidates (base rate; $|V \cap U| / |U|$, where U is the design universe and V is the viable subset).
- $\alpha$: Sensitivity (true-positive rate).
- $\beta$: False-positive rate.
- $a \in [0,1]$: Allocation to natural channel ($1–a$ to synthetic).
- $\delta \geq 0$: Data-dependency parameter (lift from natural priors to synthetic PPV).
- $\varepsilon_S(\Sigma, a) = \varepsilon_S^0(\Sigma)\,[1 + \delta a]$: Synthetic PPV with prior lift.

- $\mathscr{R}(a, \Sigma)$: Return per cost (novelty per unit effort) = $(dN/d\Sigma) / [a\ \Lambda_N\ c_N + (1–a)\ \Lambda_S\ c_S]$.

Proofs assume standard probabilistic tools (e.g., survival analysis for finitude, base-rate calculus for dilution) and economic optimization (equal-marginal rule). Where needed, I reference appendix equations and provide explicit limits/derivatives.

**P1: Allocation Gate Statement:**

At a decision horizon $\Sigma_{ref}$, if $\varepsilon_S^0(\Sigma_{ref}) < (c_S / c_N)\ \varepsilon_N / (1 + \delta\ a)$, programs do not shift to S; they invest in N or reduce β until above the gate. Proof (Derived from Appendix C: Data-dependent hybrid and equal-marginal gate):This prediction follows from the equal-marginal rule for optimizing allocation a in the hybrid model, ensuring marginal returns from natural (N) and synthetic (S) channels are balanced.

1. Viable-Discovery Rate: The hybrid rate is $dN/d\Sigma(a, \Sigma) = a\ \Lambda_N\ \varepsilon_N + (1 – a)\ \Lambda_S(\Sigma)\ \varepsilon_S(\Sigma, a)$, where $\varepsilon_S(\Sigma, a) = \varepsilon_S^0(\Sigma)\ [1 + \delta\ a]$ captures the prior lift.
2. Return per Cost: $\mathscr{R}(a, \Sigma) = [dN/d\Sigma(a, \Sigma)] / [a\ \Lambda_N\ c_N + (1 – a)\ \Lambda_S\ c_S]$. To maximize $\mathscr{R}$, take the partial derivative with respect to a and set to zero (first-order condition for interior optimum).
    - Let Rate = $dN/d\Sigma = a\ \Lambda_N\ \varepsilon_N + (1 – a)\ \Lambda_S\ \varepsilon_S^0\ (1 + \delta\ a)$.
    - Cost = $a\ \Lambda_N\ c_N + (1 – a)\ \Lambda_S\ c_S$.
    - $\mathscr{R}$ = Rate / Cost.
    - $\partial\mathscr{R}/\partial a = [\ (\partial \text{Rate}/\partial a)\ \text{Cost} – \text{Rate}\ (\partial \text{Cost}/\partial a)\ ] / \text{Cost}^2 = 0 \Rightarrow (\partial \text{Rate}/\partial a)\ \text{Cost} = \text{Rate}\ (\partial \text{Cost}/\partial a)$.
3. Marginal Rates:

- $\partial$Rate/$\partial$a = $\Lambda_N$ $\varepsilon_N$ – $\Lambda_S$ $\varepsilon_S^0$ (1 + δ a) + (1 – a) $\Lambda_S$ $\varepsilon_S^0$ δ.
- Correct expansion: Rate = a $\Lambda_N$ $\varepsilon_N$ + (1 – a) $\Lambda_S$ $\varepsilon_S^0$ + (1 – a) $\Lambda_S$ $\varepsilon_S^0$ δ a.
- Actually: (1 – a) $\varepsilon_S^0$ (1 + δ a) = $\varepsilon_S^0$ (1 – a + δ a – δ $a^2$).
- But for derivative: $\partial/\partial$a [a $\Lambda_N$ $\varepsilon_N$ + (1 – a) $\Lambda_S$ $\varepsilon_S^0$ (1 + δ a)] = $\Lambda_N$ $\varepsilon_N$ + (1 – a) $\Lambda_S$ $\varepsilon_S^0$ δ – $\Lambda_S$ $\varepsilon_S^0$ (1 + δ a).
- Simplify: $\Lambda_N$ $\varepsilon_N$ – $\Lambda_S$ $\varepsilon_S^0$ (1 + δ a) + (1 – a) $\Lambda_S$ $\varepsilon_S^0$ δ.
- Precise: $\partial$Rate/$\partial$a = $\Lambda_N$ $\varepsilon_N$ – $\Lambda_S$ $\varepsilon_S$(Σ, a) + $\Lambda_S$ $\varepsilon_S^0$ δ (1 – a) (from product rule on second term).

At equilibrium, the marginal rate from shifting to S equals that from N, adjusted for costs.

4. Breakeven Gate: For marginal shift (da > 0 from N to S), the breakeven is when marginal $\mathscr{R}$ from S ≥ marginal from N: $\Lambda_S$ $\varepsilon_S$(Σ, a) / ($\Lambda_S$ $c_S$) ≥ $\Lambda_N$ $\varepsilon_N$ / ($\Lambda_N$ $c_N$) ⇒ $\varepsilon_S$(Σ, a) ≥ ($c_S$ / $c_N$) $\varepsilon_N$.
    - Substituting $\varepsilon_S$ = $\varepsilon_S^0$ (1 + δ a): $\varepsilon_S^0$ (1 + δ a) ≥ ($c_S$ / $c_N$) $\varepsilon_N$.
    - Rearrange for gate: $\varepsilon_S^0$ < ($c_S$ / $c_N$) $\varepsilon_N$ / (1 + δ a) implies no shift (stay in N or improve S via β reduction to raise $\varepsilon_S^0$).
    - At horizon $\Sigma_{ref}$, evaluate at current a.
5. Lower Bound for a: Solving the FOC yields a ≥ [($c_S$ / $c_N$) $\varepsilon_N$ / $\varepsilon_S^0$ – 1] / δ (clipped [0,1]), as in appendix. If below gate, a = 1 (pure N).

This is falsifiable via portfolio audits at fixed $\Sigma_{ref}$.

**P2: AI Benchmark Statement:**

Exponential counter-case is passed only if measured c ≥ r and β(Σ) declines—falsifiable in prospective pipelines. Proof (Derived from Appendix B: Infinite-horizon dilution):This tests the condition to avoid dilution in synthetic space, where AI/compute enables exponential throughput but faces exponential prevalence decay.

1. Prevalence and PPV: $\pi(\Sigma) = |V \cap U| / |U| \rightarrow 0$ as $|U| \rightarrow \infty$ (dilution). PPV $\varepsilon(\Sigma) = \alpha / [\alpha + \beta (1 - \pi)/\pi] \approx (\alpha / \beta) \pi$ for small $\pi$.
2. Discovery Rate: Rate = $\Lambda(\Sigma) \varepsilon(\Sigma) \approx \Lambda(\Sigma) (\alpha / \beta) \pi(\Sigma)$.
3. Polynomial Case (Dilution Inevitable): Assume $\Lambda(\Sigma) \propto \Sigma^k$ (power-law growth), $\pi(\Sigma) \propto \Sigma^{-(k+1+\eta)}$ for $\eta > 0$ (decay faster than inverse growth).
   - Then Rate $\propto \Sigma^k * \Sigma^{-(k+1+\eta)} = \Sigma^{-(1+\eta)} \rightarrow 0$ as $\Sigma \rightarrow \infty$ (diminishing returns, per dominated convergence on bounds).
4. Exponential Counter-Case (Possible Avoidance): Assume $\Lambda(\Sigma) \approx e^{\{c \Sigma\}}$ (exponential throughput, e.g., AI scaling), $\pi(\Sigma) \approx e^{\{-r \Sigma\}}$ (exponential decay, e.g., curse of dimensionality).
   - Rate $\approx (\alpha / \beta)\, e^{\{(c - r) \Sigma\}}$.
   - For Rate $\rightarrow \infty$ (reversing dilution): $c - r > 0 \Rightarrow c > r$.
   - Additionally, if $\beta(\Sigma)$ is fixed, dilution occurs unless $c > r$; but to sustain, $\beta$ must decline (higher specificity via priors/AI), as $\varepsilon \approx (\alpha / \beta) \pi$ requires $\beta \rightarrow 0$ faster than $\pi$ decays if $\alpha < 1$.
   - Falsifiable: Measure c (throughput growth rate), r (prevalence decay rate) in pipelines; if $c < r$ or $\beta$ static/increasing, dilution persists.

This benchmarks AI: Prospective trials (e.g., screening logs) can estimate c, r, $\beta$ over $\Sigma$.

**P3: Punctuations Statement:**

Natural-channel spikes appear as transient peaks in $dN/d\Sigma$; interspike intervals $\tau_k$ lengthen over geological time. Proof (Derived from Appendix A: Natural closure):This follows from the finite first-arrival process with rare grammar expansions.

1. Finite-Class Model: M finite classes, each with hazard $\lambda_{i(\Sigma) \geq 0}$ (discovery rate), bounded/integrable. Cumulative hazard $H_i(\Sigma) = \int_0^{\Sigma} \lambda_i(s)\, ds \to \infty$.
2. Expected Discovery: $E[N(\Sigma)] = \sum_{\{i=1\}}^{M} (1 - e^{\{-H\_i(\Sigma)\}})$.
   - By dominated convergence (since $|1 - e^{\{-H\_i\}}| \leq 1$, integrable), $\lim_{\{\Sigma\to\infty\}}$ $E[N(\Sigma)] = M < \infty$ (saturation).
   - Rate $dE[N]/d\Sigma = \sum_{\{i=1\}}^{M} \lambda_i(\Sigma)\, e^{\{-H\_i(\Sigma)\}} \to 0$ (almost-sure by first-arrival theory: hazards bounded ⇒ convergence in probability, then a.s.).
3. Punctuations: Rare expansions add $\Delta M$ finite classes, resetting hazards for new ones.
   - Pre-expansion: $dN/d\Sigma \downarrow$ toward 0.
   - Post-expansion: Transient spike as new $\lambda_i$ activate, then decay resumes.
   - Intervals $\tau_k$: Over geological time, expansions rarefy (finite grammar bounds), so $\tau_k \uparrow$ (e.g., Poisson process with decreasing rate).

Falsifiable via historical/taxonomic data (e.g., COCONUT clusters).

**P4: Reverse-Eroom Regime Statement:**

If $(1 + \delta\, a) > (c_S / c_N)\, (\varepsilon_N / \varepsilon_S^0)$ (often >1.5 with defaults), a hybrid increases novelty per cost versus pure S and can raise approvals per dollar; testable in portfolio trials. Proof

(Derived from Appendix C, extending P1):This identifies the regime where hybrid ($a > 0$) outperforms pure synthetic ($a = 0$).

1. Pure S Return: $\mathscr{R}(0, \Sigma) = [\Lambda_S \varepsilon_S^0] / [\Lambda_S c_S] = \varepsilon_S^0 / c_S$.
2. Hybrid Return: $\mathscr{R}(a, \Sigma) = [a \Lambda_N \varepsilon_N + (1 - a) \Lambda_S \varepsilon_S^0 (1 + \delta a)] / [a \Lambda_N c_N + (1 - a) \Lambda_S c_S]$.
3. Dominance Condition: $\mathscr{R}(a > 0) > \mathscr{R}(0)$ when prior lift outweighs cost differences.
   - From gate: Hybrid optimal if $\varepsilon_S^0 (1 + \delta a) > (c_S / c_N) \varepsilon_N$ (rearranged: $1 + \delta a > (c_S / c_N) (\varepsilon_N / \varepsilon_S^0)$).
   - With defaults (e.g., $\delta \sim 1$, $a \sim 0.5$, $c_S / c_N \sim 1$, $\varepsilon_N / \varepsilon_S^0 \sim 1.5$), right side ~1.5, so often holds.
   - Then $a > 0$ raises $\mathscr{R}$, increasing novelty/cost → more approvals/dollar (proxy for productivity).
4. Sensitivity: $\partial a / \partial \delta > 0$ (stronger lift favors hybrid); as $\varepsilon_S^0$ ↓ (dilution), a ↑.

Testable: Randomized portfolio trials comparing hybrid vs. pure S on novelty/cost metrics.
